# Demoralization among cancer patients in mainland China: validity of the Demoralization Scale(DS)

**DOI:** 10.1101/144865

**Authors:** Lisha Deng, Ying Pang, Yi He, Yening Zhang, Richard Fielding, Lili Tang

## Abstract

Demoralization, characterized by hopelessness, helplessness, and loss of meaning and purpose, reflects existential distress. The objectives is To assess the validity of a Mainland Chinese versions of the demoralization scale (MC-DS) for using with Mainland Chinese cancer patients. In-patients sequentially recruited from a specialist tertiary-level cancer hospital in Beijing between January 2016-April 2016 completed Demoralization Scale, (DS) Patient Health Questionnaire-9 (PHQ-9), Revised Life Orientation Test (CLOT-R), Beck Hopelessness Scale (BHS), and provided sociodemographic and clinical information. We determined DS factor structure and convergent and divergent validity. 296/424 (70.0%) participants reported mean DS score=30.42(SD=13.00). EFA identified 3-factors explaining 21.4%, 17.8%, and 10.6% respectively of observed variance. Respective Cronbach Alphas were 0.88, 0.84, and 0.64 (0.90 full-scale). Convergent was shown by PHQ-9 scores correlating with Factor 2 (r=0.606), and BHS and C-LOT-R scores correlating (r=0.632,r=0.407 respectively) with Factor 1. Dichotomizing demoralization (high >30, low≦30) cross-tabulated against PHQ-9 score (mood) scores revealed 47% of patients exceeded demoralization cut-off, 60% of whom were not depressed. Using mean value±SD indicated demoralization cutoffs at <17.4 (low), 17.4-43.4 (medium) and >43.4 (high). Overall 71% met criteria for medium demoralization, and 15% for high demoralization. Sixty percent of all medium demoralization patients were not depressed, but only 5% of high demoralization patients were not depressed. The conclusion is that the Mainland Chinese Demoralization Scale is useful for detecting mild-to-moderate demoralization in cancer patients but at higher scores has poor specificity against depression.

## Introduction

Demoralization reflects existential distress [1-3], involving subjective incompetence [4], helplessness, hopelessness, worthlessness/meaninglessness, and desire for death [1, 2]. Diagnostic criteria require affective symptoms of existential distress, including hopelessness or loss of meaning and purpose in life; Attitudes reflecting pessimism, helplessness, sense of being trapped, personal failure, or lacking a worthwhile future; lacking motivation to cope differently, and; evidence of social alienation or isolation and lack of support. No major depressive or other psychiatric disorder constitutes the primary condition. Allowing for fluctuation in emotional intensity, these phenomena should persist for more than two weeks [5].

Both demoralization and depression, common in cancer [6], feature disrupted sleep and appetite, and suicidal ideation [7]. Demoralization is characterized by loss of purpose and meaning [5], depression by anhedonia [5,8]. Depressed patients struggle to experience anticipatory and consummatory pleasure, while Demoralized patients can still experience consummatory pleasure [9]. Hence, Grassi emphasized demoralization principally as existential suffering[10].

To measure demoralization syndrome a Demoralization Scale (DS) was developed [1] but scoring to classify demoralization is uncertain. Two methods have been proposed: using total score cut-off at 30 [1], or the mean value plus one standard deviation (M+1s.d.) [11].

The DS has been recently updated to improve ease of use [12] but ongoing disagreement about the divergence between demoralization and depression, and the appropriate cutoff values remain, generating methodological variability, which decreases DS diagnostic utility, so resolution is needed.

Currently, most relevant studies address English-speaking populations, though a Taiwanese Mandarin Chinese translation of the DS was validated [13]. We report a revalidation of the DS in Mainland China and compared classifications using competing methods.

## Methods

### Participants

Following ethics committee approval (2015YJ03), in-patients at a tertiary cancer center in Northern China were recruited between January 2016-April 2016 from departments specializing in alimentary, respiratory, breast, gynecological, lymphatic, genitor-urinary and musculoskeletal system malignancies. Patients 18 years or older, willing to sign informed consent and able to complete assessments were eligibile. Overall 296/424 (70.0%) of patients approached completed questionnaires.

### Assessments

Demoralization Scale (DS): Simplified characters were substituted for the traditional characters in the validated Chinese version [13]. The DS assesses status over the preceding 2 weeks. The original 24 items (α=0.94) constitute five factors: loss of meaning and purpose (α=0.87), dysphoria (α=0.85), disheartenment (α=0.89), helplessness (α=0.84), and sense of failure (α=0.71) [1]. The 5-point likert-type response scale is scored from, 0 (never), through to 4 (all the time) [11]. Five items are reverse scored, item scores are summed, higher scores reflecting greater demoralization.

Patient Health Questionnaire-9: The nine item PHQ-9 [14] measured depression over the prior two-weeks, using 4-point likert-type responses assessing symptom duration from 0 (“not at all”), to 3 (“nearly every day”) yielding a total score between 0-27. A summed score >=10 constitutes criterion for a diagnosis of depression. Scale coefficients range between 0.79-0.82.

Beck Hopelessness Scale [15]: Twenty self-report items measure hopelessness using binary (“yes”/“no”) responses giving a total scores range from 0-20. Total scores of 0-3 reflect “minimal”, 4-8 “mild”, 9-14 “moderate”, and above 14 “severe” hopelessness. Scale coefficient range between 0.85-0.93.

Chinese Life Orientation Test-Revised (C-LOT-R): The Mainland Chinese version [16] comprises six items scored on 5 point likert-type scales. The balance of scores indicates an optimistic, neutral or pessimistic orientation towards future outcomes.

Social and clinical data were gathered on age, gender, education, ethnicity, occupation, marital status and diagnosis, time since diagnosis and treatment.

### Procedure

Procedures and explanations were piloted among 30 in-patients. Subsequently eligible patients were identified from ward lists and approached for informed consent. Those agreeing were then given an introduction on self-completion of the instruments. All scales were checked and missing data were clarified with patients and completed following investigator clarifications.

### Data analysis

Data were coded and doubly entered independently by two experimenters respectively using EpiData 3.0 then checked and cleaned. After sample characteristics, frequencies and proportions descriptions, principal components analysis (PCA)-based Exploratory Factor Analysis (EFA) clarified the underlying factor structure of the DS. The number of factors was not pre-specified. Factors whose eigenvalue > 1 were extracted and retained, and with scree plot analysis informed the number of factors extracted. Parsimony guided interpretation. Independent T-test then compared demoralization and non-demoralization groups. All proportions are reported as whole numbers.

## Results

296/424 (70.0%) of eligible patients participated. Mean age was 50.3 years, (SD=12.6, range 18 to 83). Females comprised 192/296 (64.9%) of the sample. Most (263/296, 89%) reported Han ethnicity and no religion (250/296, 84%). Most (271/296, 92%) were married and educated to high school (72/296 patients, 24%), junior high school (67/296, 23%) or university (63/296 patients, 21%) levels. Participants (73/296, 25%), were retired, farmers (58, 20%), professionals (41, 14%) or held other occupations (39, 13%). Most reported monthly incomes of =<¥3000 (US$450) (94, 32%), ¥3001-¥5000 (83, 30%) and ¥5001 or more (52, 18%). Breast (82/296, 28%), and respiratory tract (66/296, 22%) tumours were most common. Mean illness duration was 2.1 years, (SD 3.2 years, range 0-19.6 years), and 1.0 years since first cancer diagnosis, (SD 1.4 years, range 0-9.7 years).

### Demoralization scale

Mean DS score was 30.42 (SD=13.00, range 0-80), close to scores reported by Kissane et al (30.82, SD=17.73) [1] and Mehnert et al (29.80, SD=10.41, range 2-61) [ 17], but higher than Mullane et al’s (19.94, SD=14.62, range 1-61) [11].

### Factor structure

A 5-factor solution provided no cleaner item-factor separation than a 4-factor solution, with four cross-loading items (items 5, 11, 22, 23). However, with 4 factors, the factor loading for item 10 was smaller than 0.4, so this was deleted, and the PCA repeated. Items 22 and 23 continued to cross-load equally on factors 1 and 2. After deleting item 22 and 23, and repeating the PCA the 4-factor solution showed no cross-factor loadings for other items. In contrast a 3-factor solution provided clean item separation without further item deletion after deleting items 10(low factor loading), 22(cross-load), 23(cross-load) and so was chosen as most parsimonious. Table 1 shows the final 3-factor solution, which explained 49.8% of observed total variance, attributable to factors 1 to 3 at 21.4%, 17.8%, and 10.6% respectively.

**Table 1.**
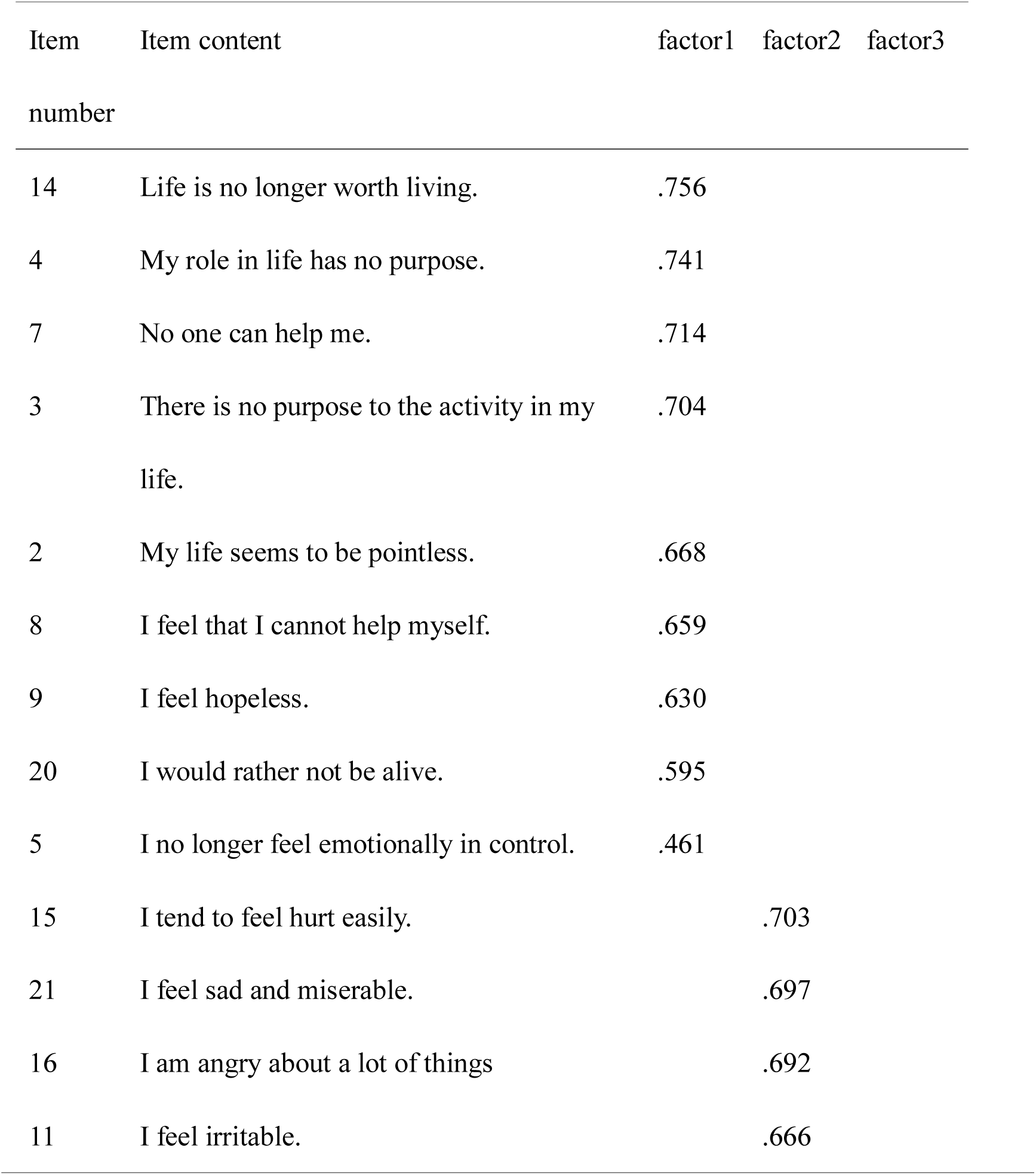
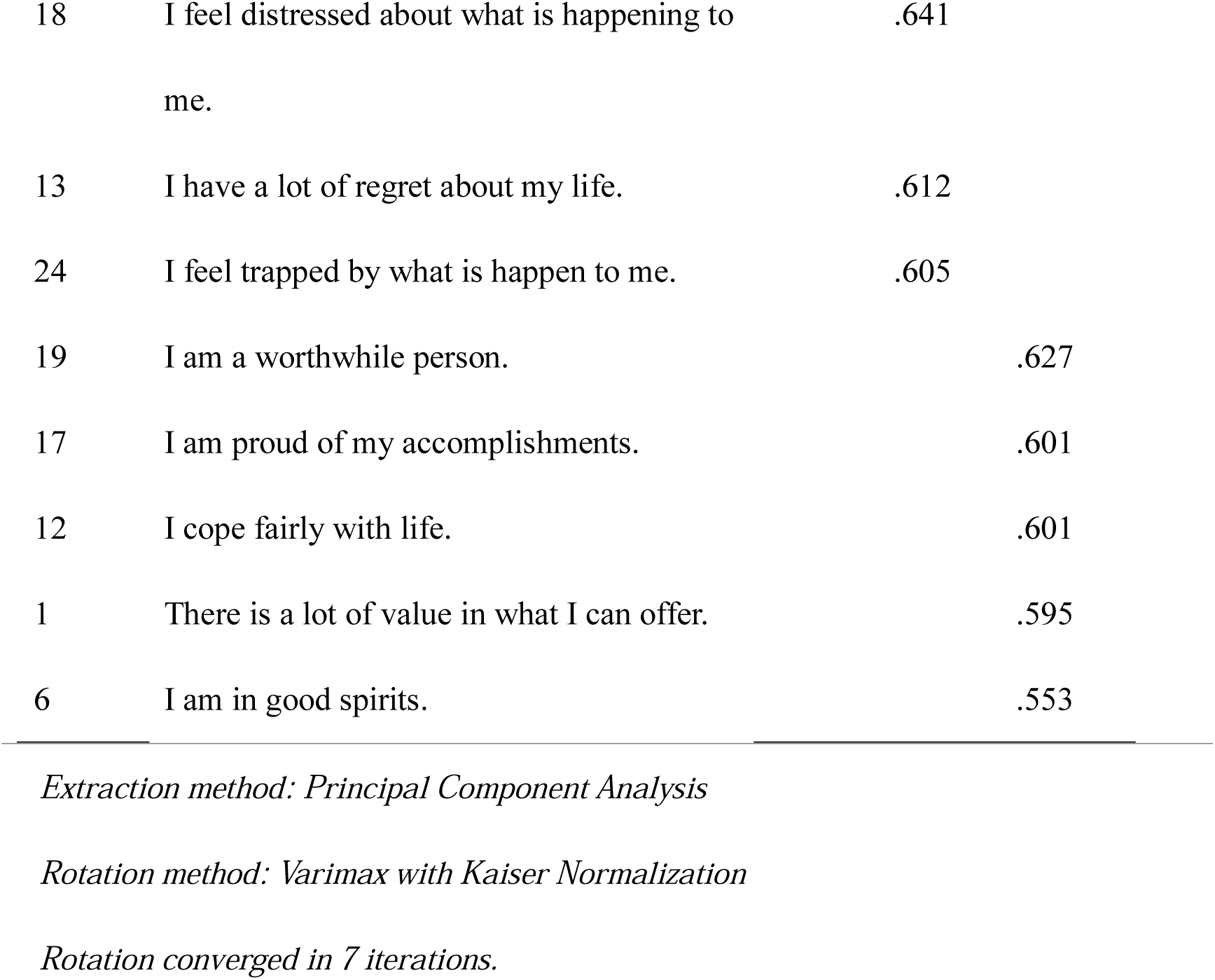
Principal components analysis of the Demoralization Scale (items 10, 22, and 23 deleted)

Factor 1 concatenated the original DS “Loss of meaning and purpose” and “Helplessness” subscales [1]. Factor 2 corresponds to the original “Dysphoria” subscale plus the “disheartenment” subscale minus item 6 (In good spirits), item 10 (Feel guilty), item 22 (Feel discouraged about life-deleted) and item 23 (Feel isolated or alone-deleted). Factor 3 corresponds to the original “Sense of failure” factor plus item 6 (In good spirits). These three factors make good intuitive sense, capturing Despondency, Distress and Self-worth, and were so named.

Cronbach’s *α* of the total demoralization scale was 0.90, and Cronbach’s *α* for factors 1 to 3 were 0.88, 0.84, and 0.64 respectively, indicating good (Factors 1 & 2) to acceptable (Factor 3) item scalability. Subscale intercorrelations were 0.606 between Factor 1 and Factor 2, 0.489 between Factor 2 and Factor 3 and 0.478 between Factor 1 and Factor 3 (Table 2).

**Table 2:** Intercorrelation among subscales of Demoralization Scale

| subscale | Factor 1 | Factor 2 | Factor 3 |
| --- | --- | --- | --- |
| Factor 1 | 1 | 0.606<sup>a</sup> | 0.478<sup>a</sup> |
| Factor 2 |  | 1 | 0.489<sup>a</sup> |
| Factor3 |  |  | 1 |

<sup>a</sup> : $p < 0.01$ for all values.

Mainland Chinese DS (MC-DS) subscales positively correlated with PHQ-9, BHS and negatively with the C-LOT-R. PHQ-9 score correlated most strongly with MC-DS Factor 2 (r= 0.606). The BHS correlated most strongly with MC-DS Factor 1 (r=0.632). The C-LOT-R correlated most strongly with MC-DS Factor 1 (r= -0.407) (Table 3).

**Table 3:** Correlations between DS subscales and PHQ-9, BHS, and CLOT.

|  | Factor 1 | Factor 2 | Factor 3 |
| --- | --- | --- | --- |
| PHQ-9 | 0.572<sup>a</sup> | 0.606<sup>a</sup> | 0.352<sup>a</sup> |
| BHS | 0.632<sup>a</sup> | 0.546<sup>a</sup> | 0.428<sup>a</sup> |
| CLOT | -0.407<sup>a</sup> | -0.404<sup>a</sup> | -0.389<sup>a</sup> |

*a: $p < 0.01$ level(2-tailed).*
*b: $p < 0.05$ level(2-tailed)*
*PHQ-9: Patient Health Questionnaire -9; BHS: Beck Hopelessness Scale; C-LOT-R:* *Mainland Chinese Revised Life Orientation Test; MCMQ Medical Coping Modes* *Questionnaire.*

Two approaches were used to compute the MC-DS demoralization cutoff values. First, dichotomization at the mean of 30 (>30 high, ≤30 low demoralization) [1], then by trichotomization using score tertiles [11]. For the latter, given our observed mean of 30.42, SD=13.00, MC-DS total score was divided into tertiles of low demoralization (<17.4), medium demoralization (17.4-43.4) and high demoralization (>43.4). Table 4 shows the resulting classifications. When dichotomized at mean score 47% of the sample met the demoralization criterion. In contrast, trichotomized cutoff values classified 71% of all patients as having medium demoralization and 15% of all patients as having high demoralization.

**Table 4.**
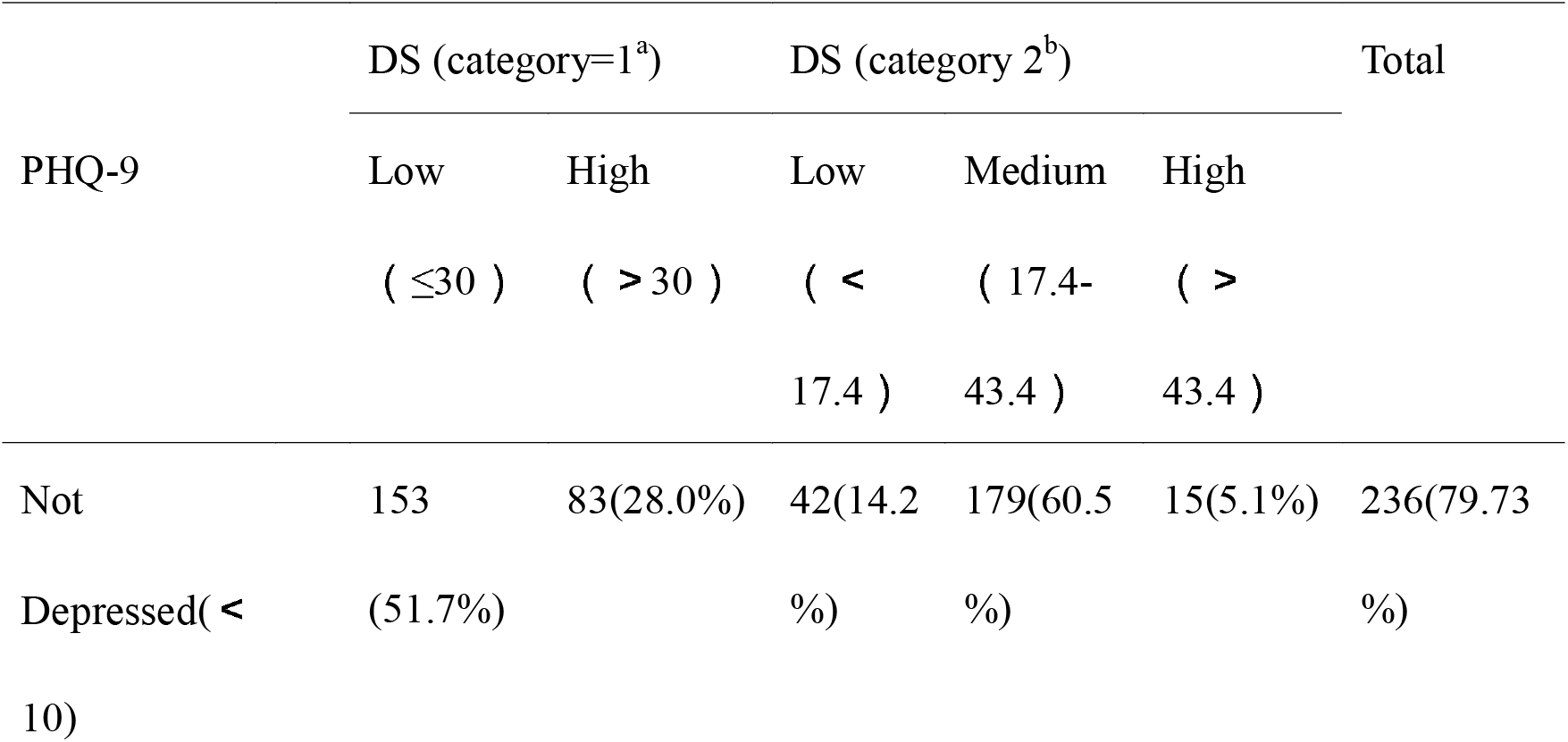
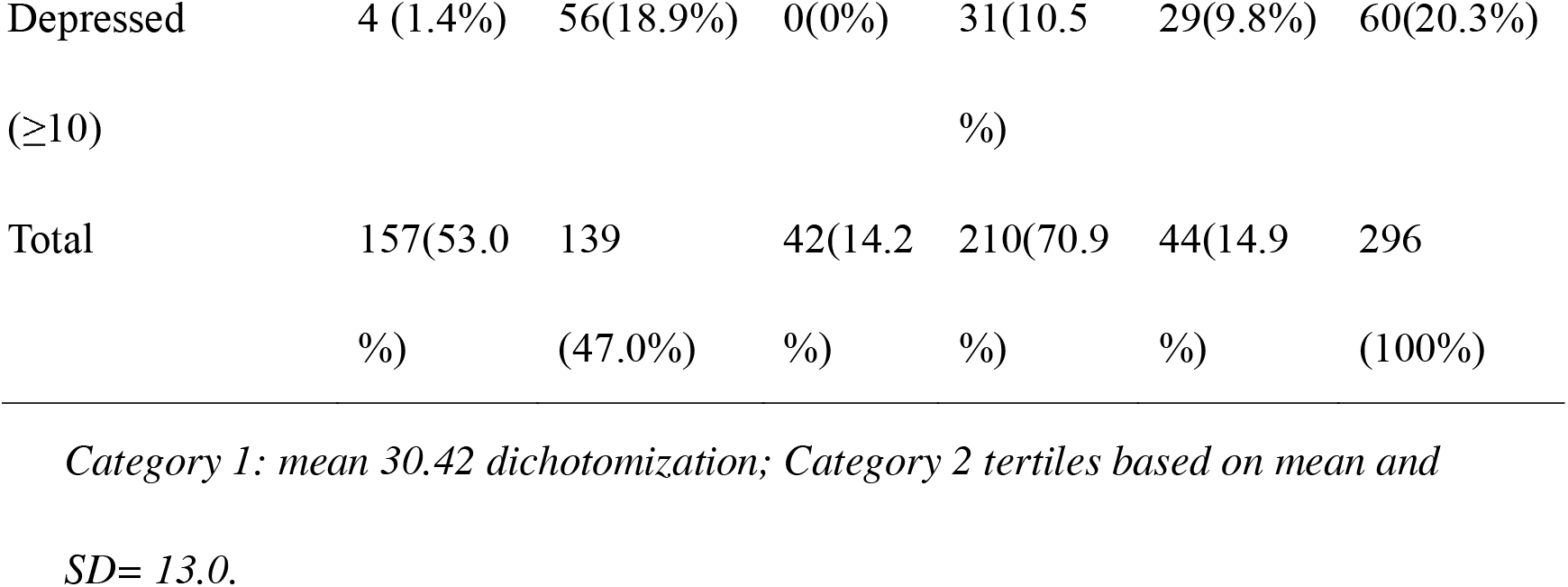
Cross-tabulation of Demoralization Scale (DS) and Patient Health Questionnaire -9(PHQ-9)

The ability of the MC-DS to discriminate between demoralization and depression determines its utility. Of 296 respondents, 60/296 (20%) achieved PHQ-9 scores indicating likely depression. Of these 56 (93%) also met the demoralization criterion when dichotomizing; when trichotomizing all 60 (100%) met medium- or high-demoralization criteria. Among those trichotomized as low demoralization only 1-in-40 were likely depressed (false negative). Conversely, dichotomization identified 47% of participants as demoralized, 19% of who were also depressed but, of the 53% not classified as demoralized, only 1.4% were depressed. This suggests that depressed individuals in this population will almost always attract a Demoralization diagnosis using these criteria and instrument.

Of respondents identified as demoralized, approximately 1-in-5 were also depressed, irrespective of the classification method used. Using trichotomization, 15% of medium- and 66% of high-Demoralized cancer patients were also likely depressed, but none of the 42 respondents in the low demoralization category were depressed. Trichotomization shows that most demoralized patients fall within 1 S.D. either side of the sample mean, whereas most depressed patients score above the sample mean on the DS. Those scoring higher than DS mean+1 S.D have a 2-in-3 chance of being depressed, while those scoring lower than DS mean-1 S.D. will not be depressed.

## Discussion

Among 296 Chinese cancer in-patients the adapted MC-DS generated a mean score=30.42, SD=13.00, similar to reported Australian [1] German samples [17], but higher than an Irish sample [11].

Kissane et al’s [1] 5-factor solution adopted for Lee et al’s Taiwanese validation [2] did not readily fit the Mainland Chinese sample reported herein. Several items cross-loaded on multiple factors and another (item 10, “feeling guilty”) was ejected for low loading. After testing 5-, 4-, and 3-factor solutions, and requiring deletion of three items a 3-factor model proved most parsimonious. Two deleted items (feeling discouraged/alone) do not discriminate adequately in the highly group-oriented Chinese culture as Confucian moral codes dictate the sick be cared for and protected by their families. Families support patients and often diagnosis is withheld to maintain hope, reflecting beliefs that loss of hope precipitates rapid deterioration and premature death.

This 3-factor solution accounted for 49.8% of observed score variance and compares sensibly with the original 5-factor structure [1]. Factor 1 concatenated Kissane et al’s “helplessness” and “loss of meaning and purpose” factors, into a factor named “Despondency” that captures the sense of this item cluster. Our Factor 2, labelled “Distress” captures Kissane et al’s “dysphoria and disheartenment” and elements of “sense of failure” factors reflecting hurt, anger, frustration and regret. The remainder of “sense of failure” loaded on our Despondency factor. Our final factor 3 clearly captures items addressing perceived “Self-worth” and is labeled as such. These three factors we feel provide a good representation of demoralization within a different cultural context. Our Despondency factor did not differentiate what Kissane et al [1] called “loss of meaning and purpose” from “helplessness”, suggesting a core element of demoralization. Similarly our Distress factor did not differentiate “dysphoria” from “disheartenment” suggesting a degree of conceptual commonality.

Mood scores (PHQ-9) positively and strongly correlated with all three factors, less so with self-worth and more so with Despondency, whereas hopelessness scores (BHS), as expected, also correlated most with factor 1, Despondency.

Both existing classification cut-offs indicate that as DS scores increase above the mean, cancer patients are increasingly likely to be depressed (as well as demoralized) and the utility of the scale becomes more questionable. The discriminant validity (know case method) of the MC-DS declines as scores increase indicating the instrument is most useful for excluding demoralization in cancer patients scoring below the original cut-off [1].

When using the recommended dichotomization scoring method [1], 47% patients were “demoralized”, while of patients not depressed on PHQ-9 scores 28% scored high demoralization. That substantially exceeds the 7%-14% identified by Kissane et al [1] and the 19.6% identified by Mehnert et al [17]. In contrast, with the trichotomization method [11] a proportion, 71% of the present sample, comparable to the 72.1% reported by Mullane et al [11] were classified as having medium demoralization, while 15% met the criterion for high demoralization, approximating the 13.4% reported by Mullane et al [10]. A higher proportion of 60% met the criterion for medium demoralization without depression, slightly higher than the 51.5% reported by Mullane et al [11], and 5% of these Chinese patients identical to Mullane et al’s 5.2% met the criteria for high demoralization without depression. The similarity of the classification proportions in this Chinese and Mullane’s Irish sample are noteworthy. This suggests the trichotomization scoring method may be more robust to sample variability. Clarification of this point is needed.

Our results support concurrent validity, but not divergent validity of the DS. At higher values, the DS poorly differentiates demoralization from depression. This is consistent with the results of Mullane, et al [11] and Mangelli et al [18], but not Kissane et at [1] Mehnert, et al [17] and Hung et al [13]. The scale in its current form does not readily differentiate demoralization from depression in this population, except at low levels of demoralization, and below scores of mean-1s.d. where demoralized patients are unlikely to be depressed.

This study has several limitations including, patient recruitment from only one cancer center, though this should not make a large difference in this type of study. Using only inpatients may bias the apparent prevalence of demoralization reported in this study. However, again, determining the absolute prevalence of demoralization was not the purpose of this study and so this, in fact, is a strength because it increases the power of the study to differentiate between demoralized and depressed.

Demoralization was measured in cancer patients, but this says nothing about whether the demoralization scale would fit other populations in China, for example those with schizophrenia, heart failure or end-stage renal disease. Otherwise, the sample size was adequate for the number of scale items and the analysis was performed as an EFA and not a CFA, thereby avoiding the assumptions underpinning the latter.

## Acknowledgments

We thank the participation of patients in Beijing Cancer Hospital.

